# Area compressibility moduli of the monolayer leaflets of asymmetric bilayers from simulations

**DOI:** 10.1101/689679

**Authors:** J. F. Nagle

**Affiliations:** Carnegie Mellon University

## Abstract

Extraction from simulations of the area compressibility moduli of the monolayers in a bilayer is considered theoretically. A statistical mechanical derivation shows that the bilayer modulus is the sum of the two monolayer moduli, as is often supposed, but contrary to a recent study. Seemingly plausible assumptions regarding fluctuations are tested rigorously. Prospects for future research are discussed.

**Significance:** It is important to describe the properties of both leaflets of generally asymmetric Biomembranes. One such property is the area compressibility modulus. This MS rigorously establishes the fundamental theory that corrects a recent BJ paper. The theory is straightforward but substantial enough that it was not readily apparent why the previous theory was incorrect. This is why this MS should be considered a new paper and not just a comment. Another reason is that this MS points to an alternative method, used only once previously, for extracting the leaflet area compressibility modulus from simulations.

## Introduction

Biomembranes are generally asymmetric, so increasing attention has been paid to creating asymmetric model systems, both in vitro and in silico. Then, it is appropriate to consider separately the physical properties of each of the two monolayers in asymmetric lipid bilayers. Separating some of those properties experimentally is difficult, so it is appropriate to turn to simulations. Those simulations that agree with experiment for all the properties that experiment can measure can then be considered for extracting properties that experiments do not measure (*1, 2*). The property of interest in this paper is the area compressibility modulus. There are two well-known methods of extracting the bilayer modulus from simulations. This paper focuses on the extraction of the individual monolayer moduli.

An area compressibility modulus k is generally defined as

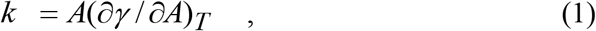

where *A* is the area and γ is the surface tension. This modulus is essentially a spring constant. Assuming that there is negligible coupling between the two monolayers, *j*=1 and 2, each monolayer can be thought of as analogous to a spring with modulus *k*_*j*_ and the bilayer would then be two springs of equal length in parallel subject. The forces on the springs would be *F*_j_ = *k*_j_ *x* and the force on the two springs would be *F*_12_ = *F*_1_ + *F*_2_ = (*k*_1_ + *k*_2_) *x* which is then identified as *k*_12_ *x*. It would then follow by analogy from elementary mechanics of springs that the modulus *k*_*12*_ for a bilayer is the sum of the monolayer moduli would be

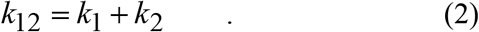

In contrast to Eq. 2, a recent paper derived a rather different equation (*3*).

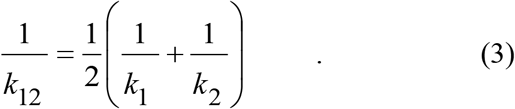

This equation has two highly unusual features. The first comes from applying it to a symmetrical bilayer. Then the two monolayer moduli must be equal, *k*_*1*_=*k*_*2*_, so Eq. 3 requires that each monolayer modulus must equal the bilayer modulus *k*_*12*_. This unusual feature was specifically noted and a rationalization was provided (*3*). The second unusual feature comes from considering a bilayer that is highly asymmetric, for example, monolayer one might consist of gel phase DPPE at room temperature and monolayer two might consist of highly fluid DOPC. In the limit when *k*_*1*_ is very much larger than *k*_*2*_, Eq.3 predicts that the bilayer modulus *k*_*12*_ is only twice the smaller monolayer modulus *k*_*2*_. This violates the definition in Eq. 1 because the tension γ_1_ to change the area of monolayer 1 should be enormous compared to the tension γ_2_ to effect the same change to the area of monolayer 2. A macroscopic analogy would be to construct a bilayer consisting of a sheet of rubber on a sheet of steel and claim that the area compressibility is unrelated to that of the steel. As the derivation provided for Eq. 3 has gaps and makes unproven assumptions (*3*), it is appropriate to return to basics.

After laying the statistical mechanical foundation in the Methods Section, the current paper provides a rigorous derivation of Eq. 2 in the first Results subsection. The second Results subsection reveals exactly which assumptions employed in (*3*) are incorrect for the case of uncoupled monolayers considered there. It also allows consideration of features not considered theoretically (*3*) that would nevertheless affect that method of analyzing simulations. The Discussion assesses the prospects for applying the small patch method of (*3*) and attention is called to a different simulation method that would not be subject to the same artifacts.

## Methods

### The Theoretical System

Consider a bilayer with fluctuating area A and average area *<A>*=*A*_*0*_. The monolayer fluctuating areas *A*_*1*_ and *A*_*2*_ are necessarily constrained to be equal to the bilayer fluctuating area, *A*_*1*_ = *A*_*2*_ = *A*. The simulation method proposed in (*3*) analyses the fluctuating areas of small portions of each monolayer *j* with fluctuating areas 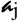. For convenience, we will set the average small areas 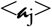 on both monolayers to be the same value 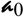. Of course, the fluctuating areas 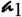 and 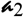 are not generally equal unless there is very strong coupling between the two monolayers. A schematic of this setup is shown in Fig. 1.

**Fig. 1.**
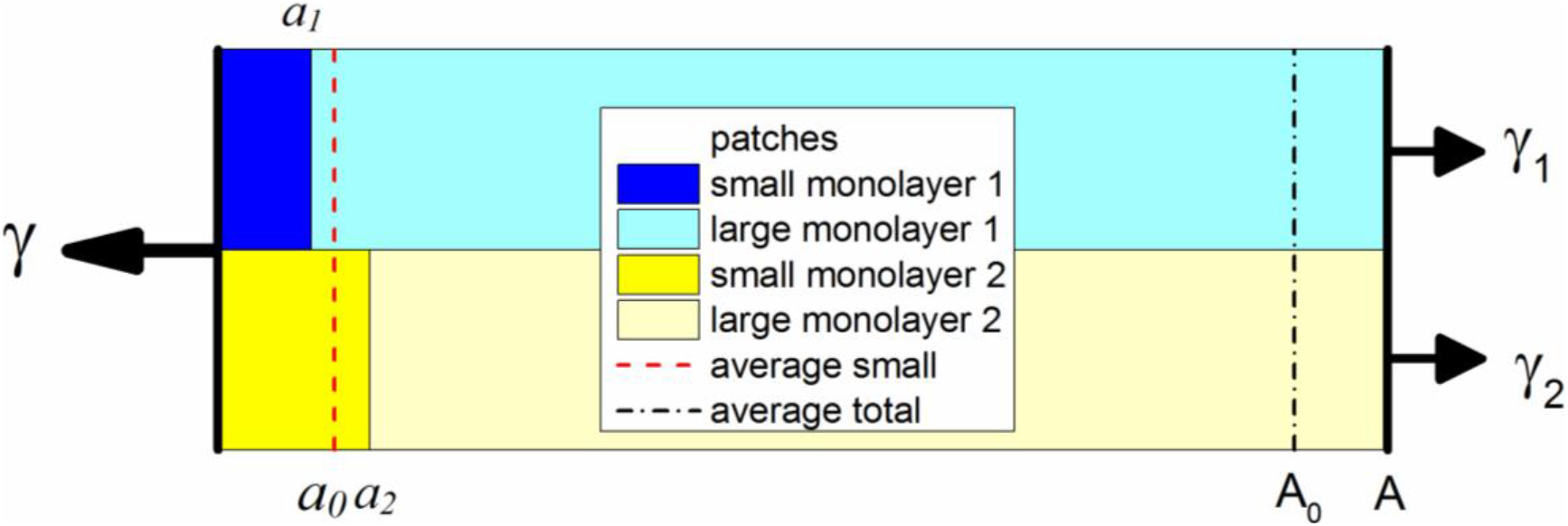
A schematic of fluctuations of the patches in two monolayers in a bilayer.

Assuming that there is also no coupling between these small fluctuating areas and between the remaining 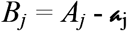 areas in each monolayer leads, via the equipartition theorem, to the monolayer moduli

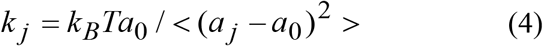

where *k*_*B*_ is Boltzmann’s constant and *k*_*B*_*T* is thermal energy. This equation is well established as one of the two main ways to obtain *k*_*12*_ when bilayer areas *A* and *A*_*0*_ replace 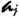 and 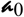 (*4, 5*). The interesting issue is how *k*_*12*_ is related to the *k*_*1*_ and *k*_*2*_ that are obtained from Eq. 4. For this, we return to the same statistical mechanics used to derive the equipartition theorem, but we now have to realize that, even if the two monolayers are uncoupled locally, there is the global constraint *A*_*1*_ = *A*_*2*_ = *A*.

The formal description of the system begins by writing the basic fluctuation energy for the small patches and also for the remaining large areas *B*_*j*_ − *B*_*0*_.

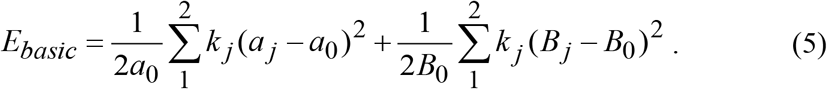

Of course, analysis of simulations would analyse many small patches in order to obtain better statistics, but there is no loss of generality in a derivation that considers small patches one at a time, each embedded in a reservoir that consists of the remaining small patches considered as a group. Each of the four terms in Eq. 5 has the conventional harmonic form for the fluctuation energy with monolayer moduli *k*_*j*_. This equation looks like it has four independent fluctuating variables, but there are only three because 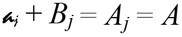. We therefore replace (*B*_*j*_ − *B*_*0*_) by 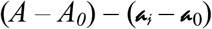 in Eq. 5. It will be convenient to condense the notation in subsequent equations by writing the three independent fluctuating variables as *x* = *A* − *A*_*0*_ and 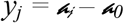. Then, Eq. 5 becomes

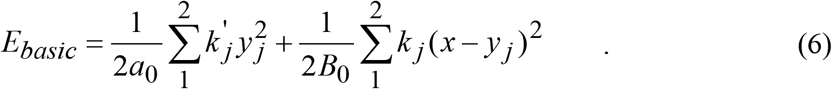

As is often the case in statistical mechanics, it is advantageous to formally distinguish nominally identical terms, such as has been done for the *k*_*j*_′ in the first term in Eq. 6, and to add terms to the basic energy as follows,

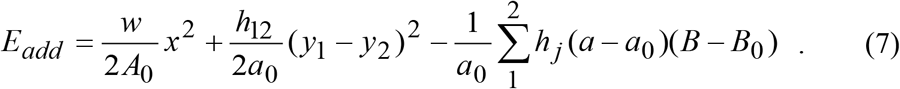

The first term in Eq. 7 is crucial because it will enable finding the relation between the bilayer *k*_*12*_ and the monolayer moduli *k*_*j*_ by taking the derivative of the partition function with respect to w and then setting w=0. The *h*_*12*_ term provides for coupling between the two monolayers. For *h*_*12*_>*0*, the coupling energy increases when the areas of the small patches are correlated. With the usual volume conservation assumption, such correlated fluctuations correspond to total bilayer thickness fluctuations (sometimes called peristaltic modes); *h*_*12*_>*0* therefore suppresses thickness fluctuations whereas *h*_*12*_<*0* enhances them. The *h*_*j*_ terms provide a kind of coupling between the small patches and the large patches in the monolayers; even more importantly, that term will enable finding the correlation functions that were previously presumed to be zero (*3*).

### Statistical Mechanical Derivation

The partition function for this system is defined as

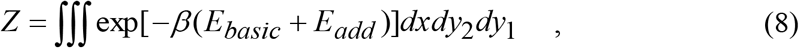

where β = *1/k*_*B*_*T*. The result of the integrations is

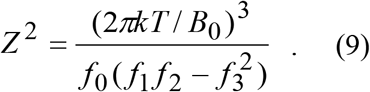

Defining *R* = (*B*_*0*_*/a*_*0*_) and *r*=(*B*_*0*_*/A*_*0*_) the *f’s* are

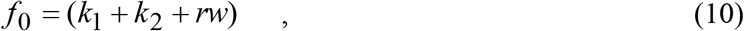

and for *j*=1 and 2

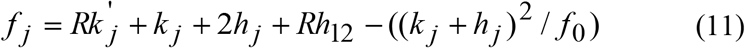

and

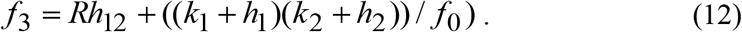

The evaluation of the partition function in Eq. 8 was performed by first grouping all the exponential factors involving *x*^*2*^ and *x*. Completion of the square in the form (a*x* − c)^2^ − c^2^ provides a Gaussian *x* integral which gives the factor *2πkT/B*_*0*_*f*_*0*_ in Eq. 9. The factors involving *y*_*1*_^*2*^ and *y*_*1*_, including those in the c^2^ factor left over from the *x* integration were then similarly treated, finally ending with a Gaussian integral over *y*_*2*_. The results of the *y*_*1*_ and *y*_*2*_ integrations together give the remaining factor in Eq. 9.

## Results

### Thermodynamic relations

Derivatives of the partition function in Eq. 9 give thermodynamic quantities of interest. First, consider the average energy

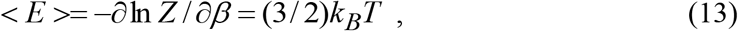

defined by the first equality in Eq. 13. The calculation using Eq. 9 gives the second equality. This recovers the usual equipartition result for three classical harmonic degrees of freedom.

The most interesting derivative is of *ln Z* with respect to the parameter *w*. By definition of the partition function in Eq. 8 and the definition of *E*_*add*_ in Eq. 7, this derivative gives the first identity in the following equation.

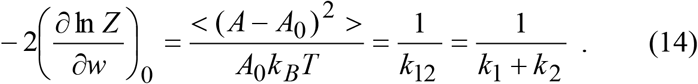

The second equality is just the identity for the bilayer modulus *k*_*12*_ as in Eq. 4. The last equality is the result of taking the derivative in Eq. 9 and then setting *w*=0 as well as *h*_*j*_=*h*_*12*_=*0*; this returns the energy to the basic terms in Eq. 6. Eq. 14 is a primary result that confirms Eq. 2 that was suggested in the Introduction by analogy to springs. This fully rigorous result proves that Eq. 3 is incorrect.

When *h*_*12*_ is non-zero, there are corrections to Eq. 14 which, however, are of order *r* and therefore vanish in the small subsystem limit 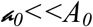. This is consistent with the infinitely strong *h*_*12*_ limit which is independently calculable because then *y*_*1*_=*y*_*2*_ is constrained and the tightly coupled monolayers reduce to a single layer with modulus *k*_*1*_+*k*_*2*_. However, the *h*_*12*_ coupling between the monolayers is far from innocuous for the interpretation of small patch fluctuations. These fluctuations are obtained by taking a derivative with respect to *k*_*j*_′ and setting *h*_*j*_=*0*=*w*, designated by 0′ in the first term in the following equation

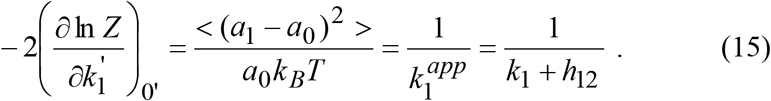

The first equality in Eq. 15 follows simply from Eqs. 7 and 8. The second equality in Eq. 15 defines the apparent monolayer modulus *k*_*1*_^*app*^ that the small patch simulation method would report. The last equality in Eq. 15 shows the result of the calculation using Eq. 9. Importantly, *k*_*1*_^*app*^ is not the true monolayer modulus *k*_*1*_, but becomes *k*_*1*_ + *h*_*12*_.

Encouragingly, one could determine *h*_*12*_ = ½ (*k*_*1*_^*app*^ + *k*_*2*_^*app*^ − *k*_*12*_) and thence obtain *k*_*1*_ and *k*_*2*_ using the final equality in Eq. 15. However, this assumes that the only coupling is between patches on opposite monolayers.

Although the *h*_*j*_ terms might appear to provide the in-plane coupling equivalent to the *h*_*12*_ out of plane term, there is a difference that makes the *h*_*j*_ terms unsatisfactory for determining *k*_*1*_ and *k*_*2*_. For either sign of *h*_*j*_ some fluctuations decrease the *h*_*j*_ energy term; that even leads to instability of the system for modest values of *h*_*j*_. However, it may be noted that these terms decrease *k*_1_^*app*^ and *k*_*12*_, but only proportional to *h*_*j*_^*2*^ and to 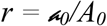. A better model for in-plane coupling might involve adding terms like 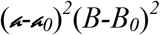 to the energy but this would introduce quartic terms which, even if calculable, would complicate an already complicated derivation of the partition function.

### Correlations

It is interesting to see exactly how plausible assumptions for correlations between the patches fail due to the *A*_*1*_ = *A*_*2*_ = *A* constraint. Let us begin with the following identity, alluded to in (*3*) that follows from *A* = ½(*A*_*1*_+*A*_*2*_).

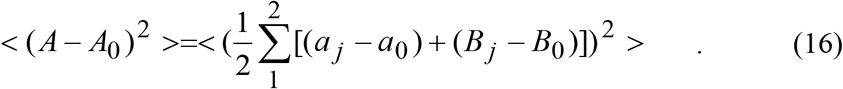

The left hand side is just *A*_*0*_*k*_*B*_*T*/*k*_*12*_. Expanding the right hand side gives

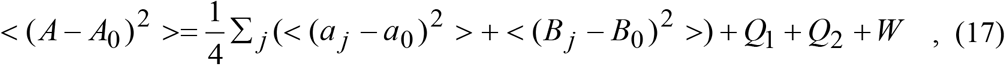

where

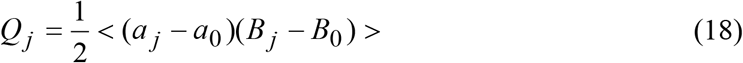

and

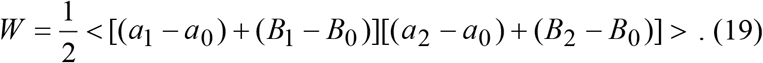

*W* was previously assumed to be zero (*3*), but it is trivially equal to ½ <(*A*−*A*_*0*_)^2^> by inspection.

Eq. 17 can now be rewritten as

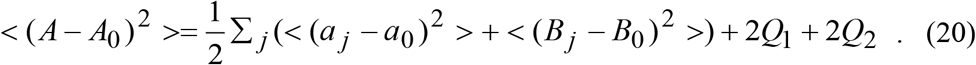

If there is no specific coupling between patches, application of Eq. 4 shows that 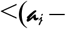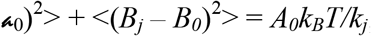, so Eq. 20 can be further rewritten as

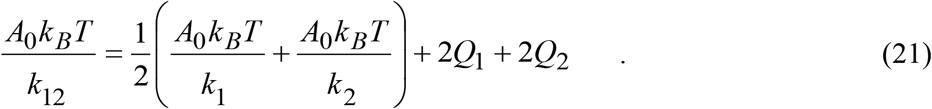

If *Q*_*1*_+*Q*_*2*_ were zero, then this would be a derivation of Eq. 3. However, the *Q*_*j*_ are straightforwardly determined to be non-zero by taking derivatives of the partition function with respect to *h*_*j*_. The result for *Q*_*1*_ is

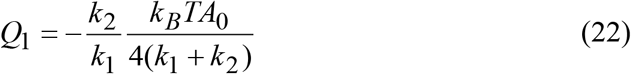

and the result for *Q*_*2*_ simply exchanges the indices 1 and 2, so *Q*_*1*_+*Q*_*2*_ is not zero. The product *Q*_*1*_*Q*_*2*_ depends only upon the sum *k*_*1*_+*k*_*2*_, but the ratio *Q*_*1*_/*Q*_*2*_=(*k*_*2*_/*k*_*1*_)^*2*^ shows that the *Q*_*j*_ have quite different values for asymmetric bilayers with larger values of *Q*_*j*_ for the softer monolayer than for the stiffer one. Finally, combining *2Q*_*j*_ in Eq. 22 with the *k*_*j*_ terms in Eq. 21 gives, for both *j*=1 and 2, the result ½*A*_*0*_*k*_*B*_*T*/*(k*_*1*_+*k*_*2*_) thereby giving

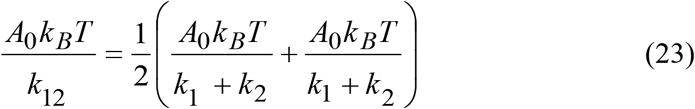

which again confirms that the bilayer modulus *k*_*12*_ is the sum of the monolayer moduli as in Eq. 2. Figure 2 plots the terms in Eq. 21 as the relative stiffness of the two monolayers varies.

**Figure 2.**
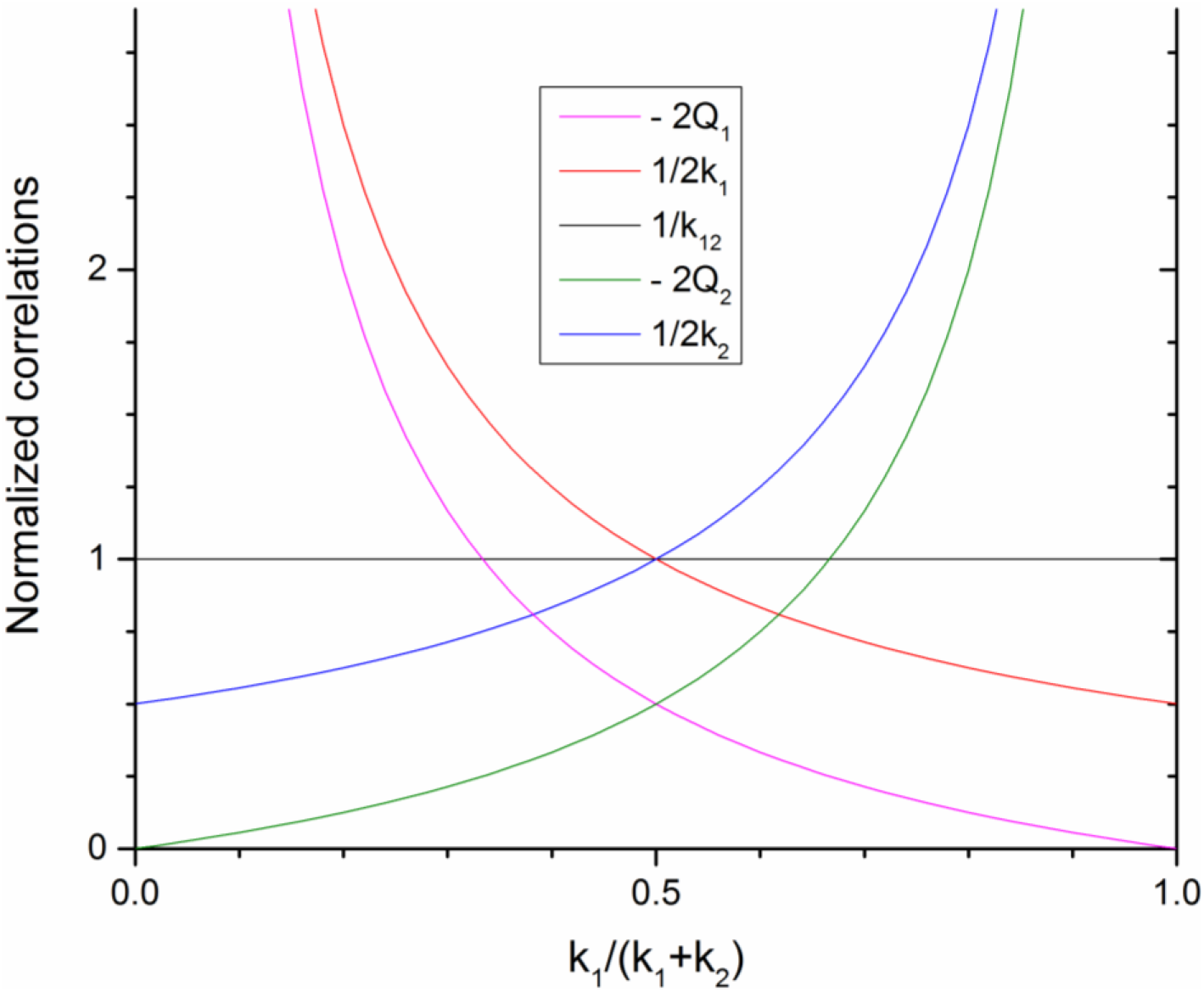
The terms in Eq. 21 are normalized to *A*_*0*_*k*_*B*_*T*/(*k*_*1*_+*k*_*2*_). The algebraic sum of the four terms on the right hand side of Eq. 21 equal 1/*k*_*12*_ = 1 since *k*_*1*_+*k*_*2*_ is normalized to 1.

It is interesting that the simple constraint that both monolayers have the same area has such a large effect on the *Q*_*j*_ and *W* correlations. Comparing to a system consisting of a single monolayer, *W* is not defined and the only defined *Q*_*1*_ is zero as one would expect from the assumption that the small patch is uncoupled from the large patch reservoir. It is also interesting to note that *W*+*Q*_*1*_+*Q*_*2*_=*0* for symmetric bilayers, but only for symmetric bilayers.

## Discussion

It is a reasonable prospect that the fluctuations in small patches will reveal differences in the monolayer moduli in asymmetric bilayers. If there is no coupling of the patches with the remainder of the bilayer, then the theory says that this analysis will give the monolayer moduli quantitatively. However, if the sum of the apparent monolayer obtained from small patches *k*_*1*_^*app*^ +*k*_*2*_^*app*^ does not equal the well determined bilayer modulus *k*_*12*_, then there must be coupling. If the coupling is only between patches on opposite monolayers, then the analysis using *h*_*12*_ allows extraction both of the coupling and the individual *k*_*j*_. Unfortunately, one could also have equality with coupling if the in-plane coupling competes with the *h*_*12*_ coupling, and we do not have a good theory for the effect of coupling within each monolayer.

The simulations previously reported (*3*) gave *k*_*1*_^*app*^ +*k*_*2*_^*app*^ = *2k*_*12*_, indicating strong coupling. However, the method employed there to convert area fluctuations to thickness fluctuations may have been flawed by the assumption that the volume was constant in a region consisting of only about half the hydrocarbon region as it was determined by the location of specific methylenes. A similar assumption has been found to be false in a recent analysis of the Poisson ratio.(*6*)

Given these problems with the small patch analysis method, it is appropriate to consider a second method that stems from the second method that has routinely been employed to obtain the bilayer modulus *k*_*12*_. This second method simply plots the area *A* versus surface tension γ to obtain the area compressibility modulus directly from its definition in Eq. 1. This method was noted in (*3*), but it was apparently not realized that it could also be used to obtain the *k*_*j*_ moduli separately. For each value of γ in the simulation one would first calculate the average lateral pressure profile Π(*z*) = −γ(*z*) of the bilayer (*5, 7, 8*). The integral of γ(*z*) along *z* across the whole bilayer is the value of γ. The idea is that one may also choose to integrate only over each monolayer separately to obtain γ_1_ and γ_2_. Then, one would calculate the separate

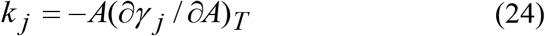

using the average <*A*> for each value of γ_j_. The only assumption in this method, as in the proposed method (*3*), is that it makes sense to separate the bilayer into two monolayers. One might argue that it does not sense to apply it to bilayers with fully interdigitated hydrocarbon chains, but it does appear to be a reasonable conceptual division for most bilayers that have only mini-interdigitation (*9*) of the monolayers near the center of the bilayer. Furthermore, one does not have to just separate *k*_*12*_ into two monolayer values. Indeed, *k*_*12*_ has already been further refined into a modulus *k*_*12*_(*z*) that varies with depth *z* for a coarse grained simulation of a symmetric bilayer (*10*). When applied to an asymmetric bilayer that method would provide an even more detailed view than just obtaining *k*_*1*_ and *k*_*2*_. As noted (*3*), this second method would require more simulations at different surface tensions and the lateral pressure profiles would probably be subject to more noise. However, this method would not be subject to the difficulties involved in the small patch method.

Finally, it may also be noted that (*3*) tackled the thorny issue of the relation between the area compressibility modulus K_A_ (=k_12_) and the bending modulus K_C_. It focused on the appropriate thickness *t* to use in

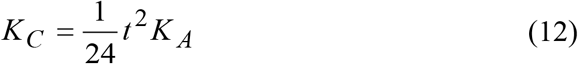

rather than on the factor of 24 that comes from the polymer brush formulation that assumes independent monolayers (*11*). The choice of the total hydrocarbon thickness for *t* has worked rather well (*2, 11*). However, the most significant exception was found when cholesterol was added to lipid bilayers and a large change in the definition of *t* was proposed (*12*). In agreement with (*3*), further analysis using reliable determinations of both *K*_*A*_ and *K*_*C*_, both from experiment and from simulations, are indeed needed to refine what effectiveness thickness *t* is appropriate to relate the mechanical properties of specific lipid bilayers.

## Conclusion

The statistical mechanical relation of the leaflet area compressibility moduli to that of the bilayer has been rigorously derived. Coupling between the two leaflets has been incorporated theoretically, and that can be addressed using the small patch simulation method. However, an alternative method is likely to be superior for obtaining more, and more reliable, information about the area compressibility of asymmetric membranes.

